# Genetic Associations with Longevity in a Calabrian Cohort: A Genome-Wide Study

**DOI:** 10.64898/2025.12.17.694890

**Authors:** Livia Beccacece, Paolina Crocco, Benedetta Torbidoni-Baldassari, Stefano Pallotti, Jie Huang, Michaël E. Belloy, Giuseppe Passarino, Giuseppina Rose, Serena Dato, Valerio Napolioni

**Author notes:** Corresponding author. Valerio Napolioni, Ph.D., School of Biosciences and Veterinary Medicine, University of Camerino, Via Gentile III da Varano, 62032, Camerino, Italy. Co-last authors (Serena Dato, Valerio Napolioni).

## Abstract

Human longevity is a complex trait shaped by genetic background and population-specific factors. Calabria, a region in Southern Italy with a high prevalence of centenarians and relative genetic isolation, is a valuable model for investigating the genetic architecture of extreme survival. Here, we performed a genome-wide association study of longevity in 705 Calabrian individuals, comparing long-lived subjects to younger controls using a mixed-model approach that accounts for relatedness and population structure. We identified 267 candidate longevity-associated variants, including 23 suggestive genomic risk loci, of which one reached genome-wide significance. Although most loci did not replicate in external datasets, one intronic variant regulating proteasome-related gene expression was confirmed by meta-analysis. Gene- and pathway-based analyses highlighted biological processes central to aging, including proteostasis, DNA repair, telomere maintenance, apoptosis, insulin signaling, inflammation, and cancer-related pathways. Notably, established longevity loci such as *APOE* and *FOXO3* were not associated, underscoring population-specific genetic effects. Overall, our findings suggest that longevity in the Calabrian population arises from a combination of unique genetic influences and conserved aging-related mechanisms, providing new insights into the molecular basis of human lifespan extension.

## 1. INTRODUCTION

Aging is an irreversible process affecting humans from sexual maturity onward and is characterized by progressive decline and loss of biological functions at the cellular, tissue, and organ levels (Gianfredi et al., 2025; Guo et al., 2022; López-Otín, Blasco, Partridge, Serrano, & Kroemer, 2013, 2023). The main features of aging are telomere shortening, genomic instability, loss of protein homeostasis, chronic inflammation, and other factors that promote the development of age-related and chronic diseases, such as cardiovascular diseases, metabolic diseases (i.e., type 2 diabetes), immune responses remodeling, cancer, and neurodegenerative diseases (Gianfredi et al., 2025; Guo et al., 2022; López-Otín et al., 2013, 2023). In this context, longevity is a complex trait that can be defined as aging in a good healthy status and marked by the survival to extreme old ages or above a specific age (Borras et al., 2020); besides, long-lived individuals usually belong to the top 10% of survivors of the respective birth cohort (van den Berg et al., 2019) and avoid or present a delayed onset of most of the age-related diseases due to maintained functional integrity (Borras et al., 2020).

Several hypotheses on the role and evolution of longevity have been proposed, and evidence suggests that longevity results from a complex interplay among genetic, epigenetic, and environmental factors (Pignolo, 2019; Sen, Shah, Nativio, & Berger, 2016). Twin and pedigree-based studies have determined that longevity is influenced by genetic factors, with estimates ranging from 10% to 30%; however, the heritability coefficient (h^2^) has yet to be accurately determined (van den Berg, Beekman, Smith, Janssens, & Slagboom, 2017; van den Berg et al., 2019). In the last decades, many genome-wide association studies have been carried out to discover the molecular mechanisms underlying this phenotypic trait and have identified several genetic loci, including *APOE* (Broer et al., 2015; Deelen et al., 2019; Gurinovich et al., 2019) and *FOXO3* (Anselmi et al., 2009; Broer et al., 2015), and many biological processes related to longevity (Smulders & Deelen, 2024).

Longevity appears to be influenced by population-specific factors, as its prevalence varies among countries and even within regions, with specific areas identified as longevity hotspots (Pignolo, 2019). Additionally, many genetic markers associated with longevity have not been consistently replicated across different populations (Smulders & Deelen, 2024), suggesting that both environmental factors and genetic variability shaped by evolutionary history play important roles (Hünemeier, 2024; Pignolo, 2019; Smulders & Deelen, 2024).

Italy is recognized as a longevity hotspot (Pignolo, 2019), with notable differences in incidence between Northern and Southern regions, likely attributable to both genetic and environmental factors, such as dietary habits, as well as sex differences (A. Montesanto, Passarino, Senatore, Carotenuto, & De Benedictis, 2008; Alberto Montesanto et al., 2017). Calabria, a region in Southern Italy (**Figure 1**), exhibits a high prevalence of centenarians (A. Montesanto et al., 2008; Alberto Montesanto et al., 2017) and has experienced multiple migratory events throughout prehistory and history, resulting in distinct genetic layers within the current Calabrian population (Bruno, Laganà, Di Lorenzo, Bruni, & Maletta, 2022). Despite these migrations, the population has remained relatively genetically stable, with low heterogeneity due to genetic isolation and high levels of inbreeding over the past three centuries (Bruno et al., 2022). Although Calabria has a high number of long-lived individuals, and several genetic association studies on candidate genes have been carried out in this population (Crocco et al., 2019; Crocco, Montesanto, Passarino, & Rose, 2016; De Rango et al., 2019), the genetic contribution to longevity in this population remains to be fully elucidated. Therefore, we aimed to investigate the polygenic basis of longevity in the Calabrian population through a genome-wide association study.

**Figure 1.**
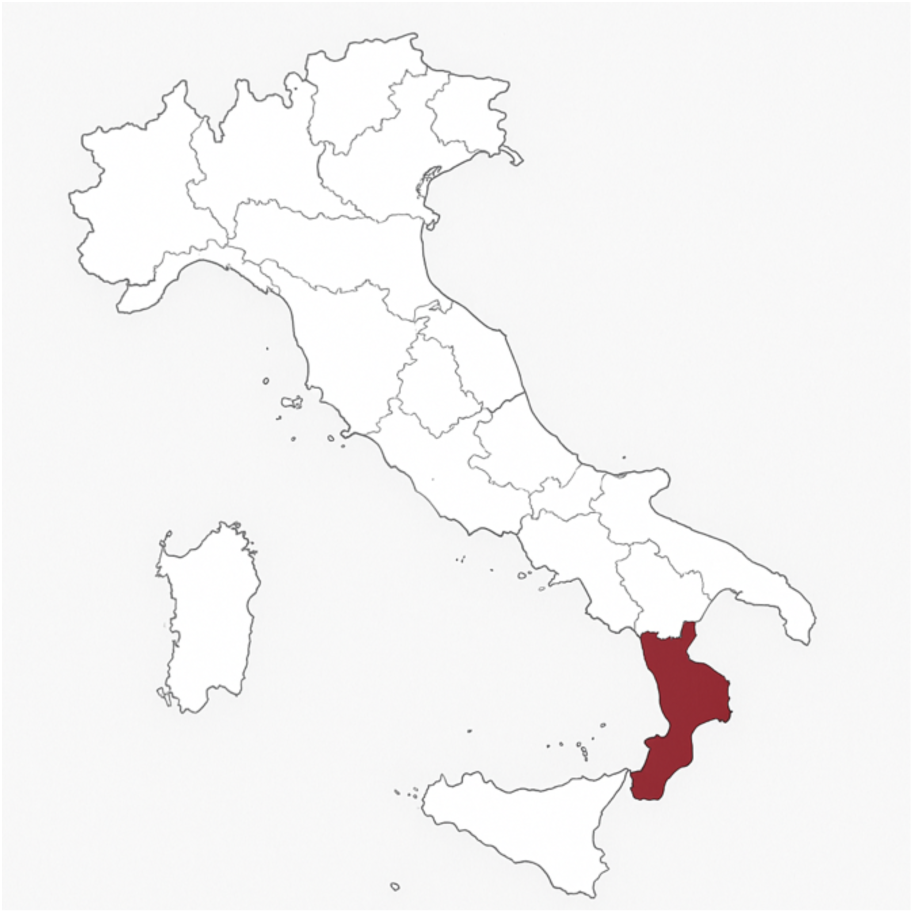
Map of Italy highlighting the Calabria region (red).

## 2. METHODS

### 2.1 Study population

The study sample comprised 723 participants (median age: 79.4 years; range: 44–107), including 310 males (median age: 77.7; range: 47–101) and 413 females (median age: 80.8; range: 44–107), recruited in Calabria (Southern Italy) through several enrolment campaigns carried out between 2002 and 2010. Written informed consent was obtained from the subjects or, where appropriate, a relative or legal representative, in accordance with institutional requirements and the Declaration of Helsinki principles. The oldest subjects (90-plus individuals) were enrolled in two European projects investigating genetic and environmental factors associated with aging: the European Challenge for Healthy Aging (ECHA) project (De Rango et al., 2008) and the GEnetics of Healthy Ageing (GEHA) project (Skytthe et al., 2011). The study protocols were approved by the Ethical Committee of the University of Calabria, respectively on 10-01-2002 for the ECHA project, and on 01-11-2004 for the GEHA projects. Younger subjects (65–85 years) were collected within the framework of several survey campaigns aimed at monitoring the health status of this population segment in Calabria (De Rango et al., 2011) (Ethical approval obtained on 09-09-2004).

The subgroup of nonagenarians (90-plus subjects; 304 individuals) included 41 individuals aged 99 years or older (32 females and nine males). The study cohort comprised related individuals. Among the 90-plus participants, 87 were singletons, 104 belonged to sibling pairs, and 3 to sibling triads. The overall cohort also included descendants of the 95+ subjects (children and grandchildren) recruited to investigate familial patterns of longevity.

A standardized questionnaire was administered to each subject, covering socio-demographic information, measures of sensory, physical, and cognitive functioning, lifestyle, morbidity (as reported by the general practitioner), psychological well-being (attitude towards life), anthropometric measurements (height, weight), and physical tests. In addition, a blood sample was collected for standard clinical hematological tests. DNA was extracted from buffy coats and stored. Vital status at baseline was traced through the official local Population Registries. An 11-year survival follow-up was performed by verifying the vital status of enrolled subjects through family contacts and official records.

To conduct a genome-wide association study of longevity, the cohort was divided into a group of long-lived individuals and a control group based on age at event/last visit. The age threshold for the assignment to the two groups was based on survival curves calculated for both males and females in the Italian population (Passarino et al., 2006). Specifically, females aged 91 years or older and males aged 88 years or older were selected as long-lived individuals. In comparison, all other individuals (females aged ≤ 90 years and males aged ≤ 87 years) were assigned to the control group. The long-lived group comprised 315 individuals (117 males and 198 females), whereas the control cohort comprised 408 individuals (195 males and 213 females).

### 2.2 Genotyping and data quality controls

After extraction, the DNA was genotyped for 654,027 SNPs using the Illumina Infinium Global Screening Array. Sex chromosomes and mitochondrial DNA were excluded from the analysis. The following filtering steps were performed on autosomal SNPs using PLINK 1.9 and PLINK 2.0 (Chang et al., 2015; Purcell et al., 2007): SNPs with a missingness rate higher than 5% (--geno 0.05) were removed, only individuals with a missingness rate lower than 5% (--mind 0.05) were retained, homozygosity levels were assessed (--het), sex checks were performed (--check-sex), and we excluded duplicated SNPs (--dup), monomorphic and multiallelic SNPs (--mac 1, --snps-only), and SNPs deviating from Hardy-Weinberg equilibrium (HWE) in controls (--hwe 1 × 10^−5^).

### 2.3 Ancestry determination, LiftOver, and imputation

In addition to the above filters, the pruning of SNPs within linkage disequilibrium (window size of 1500 bp, step of 150 bp, and r^2^ threshold of 0.1) and calculation of Identity-By-Descent (IBD) coefficients (minimum proportion of IBD equal to 0.0625) were carried out before assessing the ancestry of the population. Ancestry was assessed using SNPWeights 2.1 (Chen et al., 2013) on two external reference panels: one including Central European (CEU), Yoruba (YRI), and East Asian (ASI) populations. At the same time, the other included Ashkenazi Jews (AJW), Northwestern Europeans (NWE), and Southeastern Europeans (SEE).

The LiftOver command-line tool for Linux (Raney et al., 2024) was used to lift genomic variants from the human reference genome version GRCh37 to GRCh38. Lifted autosomes underwent pre-imputation checks to prepare the data for the genotype imputation; autosomes were checked against the HRC reference panel (McCarthy et al., 2016), determining the update of strand, position, and reference/alternative allele assignment; in addition, SNPs with differing alleles, those showing an allele-frequency difference >20%, and those not included in the reference panel were removed.

After quality checks and ancestry determination, autosomal genotypes were imputed using the TOPMed Imputation Server (Das et al., 2016) to infer millions of additional variants with high accuracy. Genotype imputation was performed using Minimac4 (Fuchsberger, Abecasis, & Hinds, 2015) against the TOPMed reference panel (version R3) (Taliun et al., 2021), applying an rsq filter of 0.3 and performing phasing.

The imputed autosomes underwent further quality checks (missingness rate, removal of duplicated and multiallelic variants, removal of SNPs deviating from HWE in controls), and we retained only SNPs with minor allele frequency (MAF) greater than 1%. This filtering was done using PLINK 1.9 and PLINK 2.0 (Chang et al., 2015; Purcell et al., 2007).

### 2.4 Association analysis

The KING-homo method (Manichaikul et al., 2010), implemented in the R package SNPRelate v. 1.40.0 (Zheng et al., 2012), was used to calculate IBD coefficients for pruned SNPs. Next, the GENESIS R package (Gogarten et al., 2019) (version 2.36.0; R software version 4.4.3) was employed to perform principal component analysis in related samples (PC-AiR) and relatedness estimation adjusted for principal components (PC-Relate). Finally, the first three principal components from PC-AiR, along with PC-Relate estimates and sex, were included as covariates in a logistic mixed-model association test to assess additive genetic effects.

The saddle point approximation (SPA) method was applied to association testing, as it provides more accurate results in the presence of binary phenotypes, case-control imbalance, and sample-relatedness (Dey, Schmidt, Abecasis, & Lee, 2017; Zhou et al., 2018).

A *p* < 5 × 10^−8^ was used as the genome-wide significance level, and *p* < 1 × 10^−5^ was used as a suggestive level of significance. Genomic control was then assessed, and the R package *qqman* v. 0.1.9 was used to visualize the results of the association test (Turner, 2018).

### 2.5 SNPs functional annotation, MAGMA analyses, and replication

The results of the association tests were annotated using the FUMA tool (Watanabe, Taskesen, van Bochoven, & Posthuma, 2017). This tool was employed to identify the functional annotation of SNPs (Wang, Li, & Hakonarson, 2010) and the deleteriousness (CADD score), and to carry out the gene-based test and the gene-set analysis through MAGMA v1.06 that uses the full distribution of SNP *p*-values (de Leeuw, Mooij, Heskes, & Posthuma, 2015); the expression quantitative trait loci (eQTLs) mapping has been performed selecting the tissue types of GTEx v8 (GTEx Consortium, 2020).

To replicate the association test, summary statistics were retrieved from a previous meta-analysis on longevity (Deelen et al., 2019); specifically, we used the summary statistics for the 90^th^ percentile of long-lived individuals of European ancestry (Deelen et al., 2019). Beta values have been meta-analyzed through a fixed-effect model using the R package *metafor* version 4.8 (Viechtbauer, 2010).

## 3. RESULTS

### 3.1 Quality checks and population structure

After quality control, 705 individuals were retained, comprising 305 males and 400 females; 18 individuals were removed owing to high missingness rates, sex-check issues, and the presence of a couple of twin sisters. Following the division applied based on the age at the last visit/event, the cohort included 299 long-lived individuals (112 males and 187 females; mean age: 97.1 years) and 406 control individuals (193 males and 213 females; mean age: 77.1 years).

All individuals were of European ancestry (81–92%), specifically of Southeastern European ancestry (66–93%), and displayed a lower proportion of African ancestry (7–10%). It is interesting to note that a proportion of European ancestry was ascribed to Ashkenazi Jews (7–33%) (**Supplementary Figures 1, 2**).

478,741 autosomal SNPs passed quality checks and were imputed, yielding 26,130,740 variants. Overall, 7,754,179 SNPs were tested for longevity and for evaluating population structure and relatedness. The estimation of relatedness identified 382 related individuals with an IBD value greater than 0.022, corresponding to 4^th^-degree relatives.

### 3.2 Association analysis and replication

Logistic mixed-model association testing, evaluating the additive genetic effect and accounting for sex and sample-relatedness, was carried out on SNPs with MAF > 1%. The genomic control inflation (λ) for this GWAS was 1.015 (**Figure 2**).

**Figure 2.**
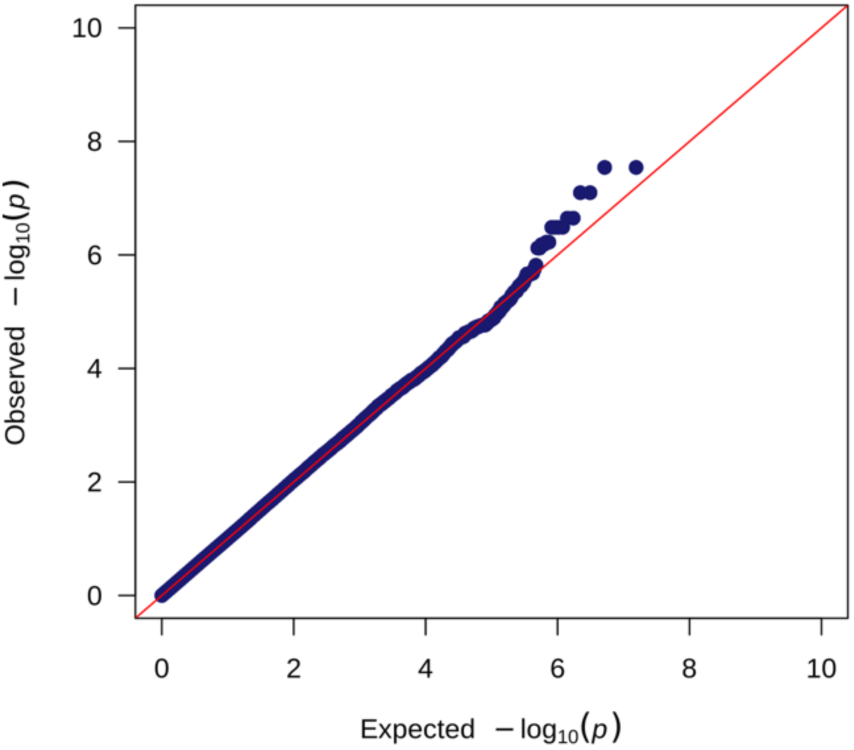
Q-Q plot of GWAS *p*-values.

The association test identified 396,583 SNPs passing the significance threshold (*p* < 0.05), of which FUMA reported 267 as GWAS candidate SNPs associated with longevity. Most of these variants are located outside coding regions (**Table S1 of Supplementary File 1**); moreover, four out of 267 SNPs have also been reported to be associated with traits unrelated to longevity (**Table S2 of Supplementary File 1**).

Among the GWAS candidate SNPs, 23 variants were identified as genomic risk loci significantly associated with longevity (*p* < 1 × 10^−5^, **Figure 3**), most of which are located within linkage disequilibrium (LD) blocks (**Table S3 of Supplementary File 1**). Only one variant, rs11186068, reached the genome-wide significance threshold (*p* = 2.87 × 10^−8^). This SNP, located within an intron of the *LINC00865* gene, which encodes a long non-coding RNA (lncRNA), is in LD with 15 other SNPs (**Table S3 of Supplementary File 1**). Another independent significant SNP, rs77148518 (*p* = 2.16 × 10^−6^), is in a separate LD block within the same genomic region (**Table S3 of Supplementary File 1)**.

**Figure 3.**
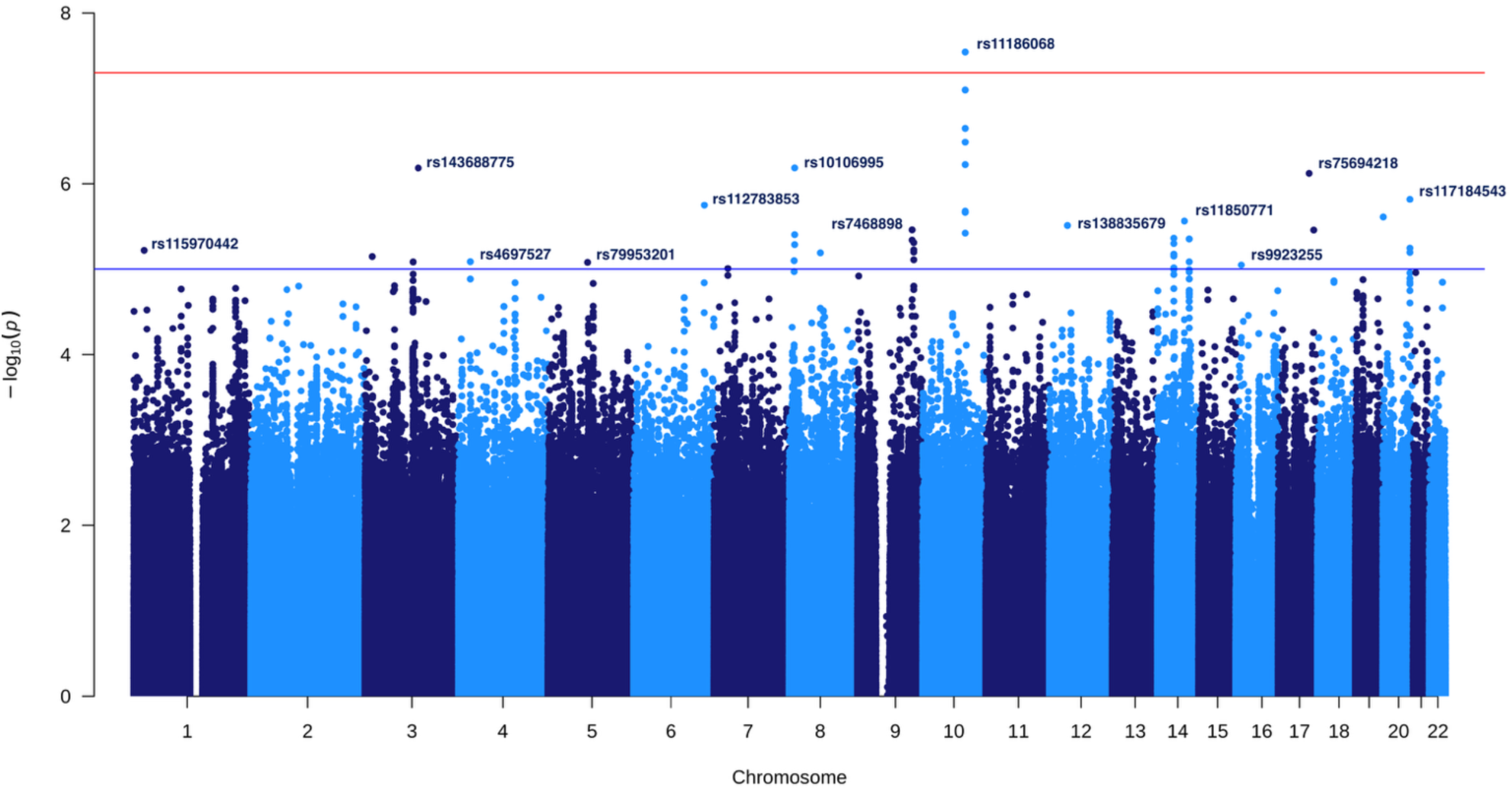
Manhattan plot of GWAS summary statistics. The red line is the genome-wide significance threshold (*p* = 5 × 10^−8^), while the blue line is the suggestive significance threshold (*p* = 1 × 10^−5^). The most significant SNPs passing the suggestive level of significance are shown.

As shown in **Table 1**, most genomic risk loci are intronic variants, primarily located in genes encoding lncRNAs and protein-coding genes, and exhibit a beneficial effect on longevity [Odds Ratio (OR) > 1] (**Table S1 of Supplementary File 1**). Notably, we did not replicate any associations for SNPs in the *APOE* and *FOXO3* genes, which have previously been reported as associated with longevity (**Table S4 of Supplementary File S2**).

**Table 1.** Genomic risk loci associated with longevity.

| Chr | SNP | Annotation | Gene | EA | Discovery cohort |  |  | Replication cohort (90<sup>th</sup>) |  |  | Meta-analysis |  |  |
| --- | --- | --- | --- | --- | --- | --- | --- | --- | --- | --- | --- | --- | --- |
|  |  |  |  |  | EAF | Beta | p-value | EAF | Beta | p-value | Beta | SE | p-value |
| 10 | rs11186068 | intronic | *LINC00865* | G | 0.033 | 1.728 | 2.872 x 10<sup>-8</sup> | 0.010 | -0.088 | 0.530 | 0.191 | 0.129 | 0.138 |
| 8 | rs10106995 | intronic | *ENSG00000300442* | T | 0.075 | 1.088 | 6.526 x 10<sup>-7</sup> | 0.070 | -0.096 | **0.044** | -0.044 | 0.047 | 0.345 |
| 3 | rs143688775 | intergenic | — | T | 0.011 | 2.506 | 6.548 x 10<sup>-7</sup> | — | — | — | — | — | — |
| 17 | rs75694218 | intergenic | — | C | 0.052 | 1.202 | 7.564 x 10<sup>-7</sup> | 0.060 | -0.046 | 0.490 | 0.036 | 0.064 | 0.577 |
| 20 | rs117184543 | non-coding exon transcript variant | *ENSG00000296630* | T | 0.138 | 0.798 | 1.520 x 10<sup>-6</sup> | 0.130 | 0.002 | 0.957 | 0.028 | 0.031 | 0.354 |
| 6 | rs112783853 | intronic | *PPP1R14C* | A | 0.015 | 2.139 | 1.784 x 10<sup>-6</sup> | 0.020 | -0.062 | 0.614 | 0.069 | 0.118 | 0.561 |
| 20 | rs113181552 | intronic | *ENSG00000286787* | T | 0.013 | 2.241 | 2.452 x 10<sup>-6</sup> | 0.040 | 0.191 | **0.002** | 0.220 | 0.062 | **3.533 x 10<sup>-3</sup>** |
| 14 | rs11850771 | 3'-UTR | *TMED8* | C | 0.215 | -0.652 | 2.736 x 10<sup>-6</sup> | 0.190 | 0.003 | 0.902 | -0.017 | 0.025 | 0.487 |
| 12 | rs117927270 | intronic | *ENSG00000258119* | C | 0.011 | 2.326 | 3.082 x 10<sup>-6</sup> | — | — | — | — | — | — |
| 9 | rs7468898 | intronic | *LOC105376234* | T | 0.491 | 0.556 | 3.466 x 10<sup>-6</sup> | 0.450 | -0.032 | 0.095 | -0.017 | 0.019 | 0.360 |
| 17 | rs11654903 | intronic | *LLGL2* | T | 0.036 | 1.419 | 3.487 x 10<sup>-6</sup> | 0.030 | -0.154 | 0.116 | -0.015 | 0.094 | 0.872 |
| 14 | rs11158017 | intronic | *SAMD4A* | G | 0.281 | 0.608 | 4.342 x 10<sup>-6</sup> | 0.290 | 0.001 | 0.954 | 0.016 | 0.021 | 0.435 |
| 14 | rs115024289 | intronic | *LINC02296* | T | 0.061 | 1.078 | 4.433 x 10<sup>-6</sup> | 0.020 | 0.001 | 0.985 | 0.103 | 0.074 | 0.161 |
| 9 | rs564752 | intronic | *BRINP1* | C | 0.199 | 0.661 | 5.954 x 10<sup>-6</sup> | 0.230 | 0.025 | 0.262 | 0.040 | 0.022 | 0.073 |
| 1 | rs115970442 | intronic | *EPHB2* | T | 0.013 | 2.211 | 6.028 x 10<sup>-6</sup> | — | — | — | — | — | — |
| 8 | rs11784121 | intronic | *RRS1-DT* | A | 0.240 | 0.622 | 6.440 x 10<sup>-6</sup> | 0.210 | -0.034 | 0.153 | -0.015 | 0.024 | 0.520 |
| 3 | rs75901608 | intronic | *SH3BP5* | C | 0.042 | 1.297 | 7.109 x 10<sup>-6</sup> | 0.060 | -0.018 | 0.672 | 0.008 | 0.042 | 0.843 |
| 8 | rs118154613 | intronic | *PRSS51* | T | 0.019 | 1.832 | 7.940 x 10<sup>-6</sup> | 0.050 | -0.008 | 0.874 | 0.017 | 0.050 | 0.738 |
| 4 | rs4697527 | intronic | *LGI2* | C | 0.033 | 1.465 | 8.157 x 10<sup>-6</sup> | 0.030 | -0.026 | 0.745 | 0.055 | 0.079 | 0.483 |
| 3 | rs595230 | intronic | *NXPE3* | A | 0.219 | -0.612 | 8.214 x 10<sup>-6</sup> | 0.210 | -0.011 | 0.634 | -0.028 | 0.023 | 0.224 |
| 5 | rs79953201 | intronic | *HAPLN1* | C | 0.021 | 1.763 | 8.309 x 10<sup>-6</sup> | — | — | — | — | — | — |
| 16 | rs9923255 | intronic | *LITAF* | T | 0.278 | -0.590 | 8.937 x 10<sup>-6</sup> | 0.310 | 0.021 | 0.343 | 0.005 | 0.022 | 0.830 |
| 7 | rs39102 | intronic | *CHN2* | T | 0.314 | 0.544 | 9.821 x 10<sup>-6</sup> | 0.330 | 0.002 | 0.933 | 0.018 | 0.021 | 0.397 |

\* **Legend.** EA: effect allele; EAF: effect allele frequency; SE: standard error of meta-analysis.

The genomic risk loci identified in our study have not been previously associated with longevity, except for two variants, rs10106995 and rs113181552, which showed a weak association with longevity in the 90^th^ percentile of long-lived individuals in the replication cohort (Deelen et al., 2019). However, only rs113181552 maintained significance after meta-analysis of beta values (**Table 1**). This intronic variant is in *ENSG00000286787*, which encodes an antisense RNA for the *PSMF1* gene (Proteasome Inhibitor Subunit 1).

Overall, 120 candidate SNPs have been identified as eQTLs affecting the gene expression in multiple tissues, of which 8 correspond to genomic risk loci, while the remaining SNPs are GWAS candidate variants (**Table S5 of Supplementary File 1**).

### 3.3 MAGMA gene-based test and gene-set analysis

A total of 18,688 protein-coding genes were tested in the MAGMA gene-based analysis. The analysis identified 995 genes associated with longevity (*p* < 0.05), but none passed the Bonferroni-corrected significance threshold (*p* = 2.676 × 10^−6^), as shown in **Figure 4**. The most significant gene was *TRMT10C* (*p* = 2.777 × 10^−5^).

**Figure 4.**
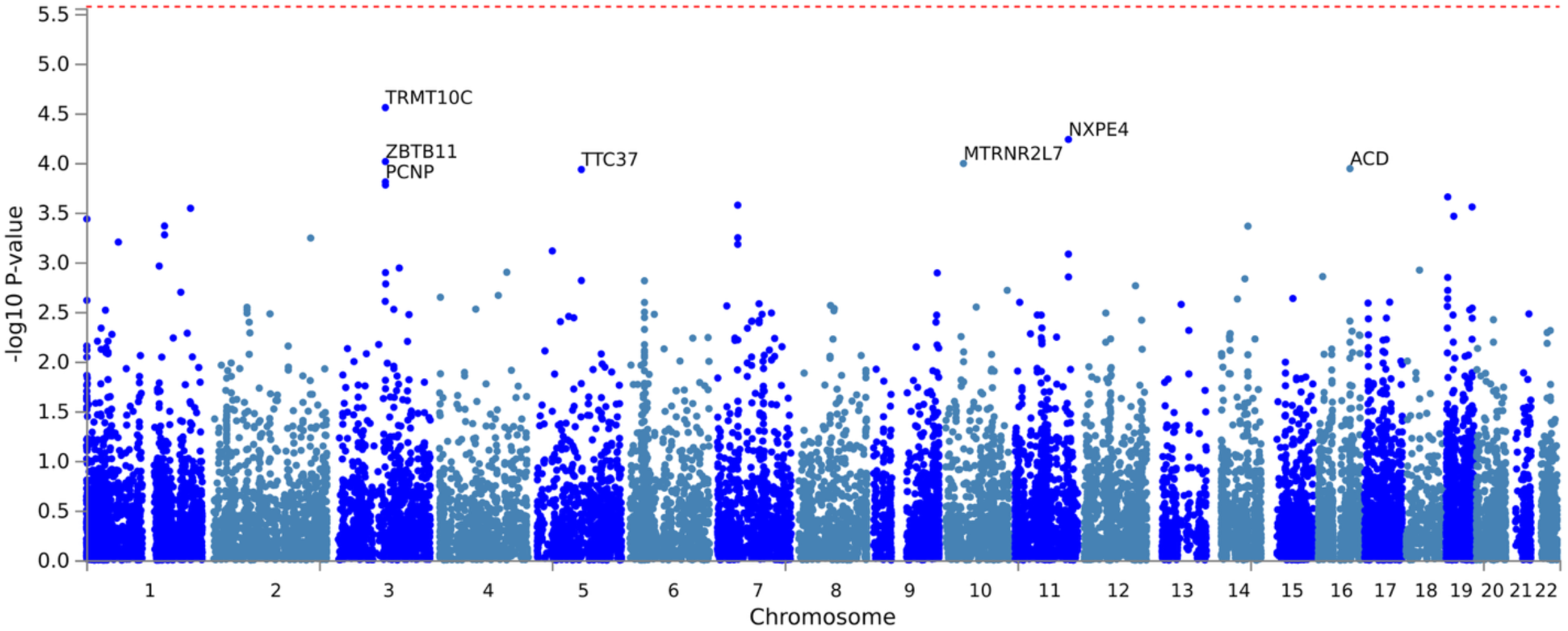
Manhattan plot of gene-based test. The red dashed line is the significance threshold of the Bonferroni correction (*p* = 2.676 × 10^−6^).

The complete list of all analyzed genes is provided in **Table S6 of Supplementary File 1**.

The MAGMA gene-set analysis identified 878 pathways nominally associated with longevity (*p* < 0.05), but none of these passed the Bonferroni correction threshold (**Table S7 of Supplementary File 1**). The ten most significant pathways are presented in **Table 2**.

**Table 2.** Most 10 significant gene-sets associated with longevity.

| Pathway | N genes | Beta | SE | *p*-value | *p*<sub>BON</sub> |
| --- | --- | --- | --- | --- | --- |
| GOMF: Insulin receptor substrate binding | 13 | 0.977 | 0.233 | 1.409 x 10<sup>-5</sup> | 0.240 |
| YU: MYC targets up | 43 | 0.396 | 0.110 | 1.681 x 10<sup>-5</sup> | 1 |
| GOBP: Positive regulation of acute inflammatory response | 26 | 0.628 | 0.176 | 1.847 x 10<sup>-4</sup> | 1 |
| INAMURA: Lung cancer SCC subtypes up | 15 | 0.782 | 0.220 | 1.859 x 10<sup>-4</sup> | 1 |
| SHEDDEN: Lung cancer poor survival A6 | 436 | 0.134 | 0.038 | 2.283 x 10<sup>-4</sup> | 1 |
| GOBP: Antibacterial humoral response | 51 | 0.445 | 0.129 | 2.859 x 10<sup>-4</sup> | 1 |
| GOMF: Preribosome binding | 8 | 1.063 | 0.313 | 3.407 x 10<sup>-4</sup> | 1 |
| HOLLERN: Solid nodular breast tumor up | 9 | 1.073 | 0.322 | 4.324 x 10<sup>-4</sup> | 1 |
| GOMF: Calcium monoatomic cation antiporter activity | 11 | 0.782 | 0.241 | 5.924 x 10<sup>-4</sup> | 1 |
| GOCC: Clathrin-sculpted monoamine transport vesicle | 5 | 1.126 | 0.348 | 6.104 x 10<sup>-4</sup> | 1 |

**\*Legend:** GOMF: Gene Ontology Molecular Function; GOBP: Gene Ontology Biological Process; GOCC: Gene Ontology Cellular Component; YU:

The most significant pathway identified was insulin receptor substrate binding (GOMF, *p* = 1.409 × 10^−5^), which involves the binding of insulin receptor substrates (IRSs) that mediate insulin signaling (Johnston, Pirola, & Van Obberghen, 2003). Overall, the top ten pathways are primarily involved in cancer, inflammation and immune response, calcium antiport, and monoamine transport.

## 4. DISCUSSION

To date, this study represents the most extensive genome-wide association study aimed at uncovering the molecular bases of longevity in the Calabrian population. Calabria region in Southern Italy is characterized by a high prevalence of centenarians (A. Montesanto et al., 2008; Alberto Montesanto et al., 2017) and genetic stratification resulting from ancient migration flows and genetic isolation over the past three centuries (Bruno et al., 2022). All Calabrian individuals included in this study mainly showed Southeastern European ancestry, with a smaller proportion of African ancestry (**Supplementary Figures 1, 2**), consistent with known migration patterns in Southern Italy (Raveane et al., 2019; Sarno et al., 2017). Interestingly, they also showed variable levels of European ancestry attributable to Ashkenazi Jews (**Supplementary Figures 1, 2**). The Ashkenazi population is known for genetic isolation, and its longevity is influenced by rare coding variants (Carmi et al., 2014; Lin et al., 2021). Although it has received some European input, said population has a substantial Middle Eastern origin (Livni & Skorecki, 2025; Santachiara Benerecetti et al., 1993). Thus, the similarity between Middle Easterners and Jews may reflect a Middle Eastern genetic component in Calabrians, which has previously been described as one of the major components of Southern Italians (Semino et al., 2000).

The identification of new genomic risk loci, along with the absence of associations for loci widely linked to longevity in other populations (e.g., *APOE* and *FOXO3*), suggests that population-specific effects may influence longevity in the Calabria region (Caruso et al., 2022). Nevertheless, this study also supports the involvement of molecular mechanisms previously implicated in longevity. Most of the genomic risk loci identified are intronic variants, several of which are located within genes encoding long non-coding RNAs (lncRNAs). Notably, lncRNAs have been associated with longevity, as some promote anti-senescence mechanisms by acting as genomic stabilizers, enhancing DNA repair pathways, and maintaining protein homeostasis (Huang et al., 2025). Among the identified genomic risk loci, only the rs11186068 polymorphism reached GWAS significance (*p* = 2.87 × 10^−8^), but it did not replicate in the validation cohort (**Table 1**). This variant has not been previously associated with any phenotype and is located within an intron of *LINC00865*. This gene encodes an lncRNA and, although its role is not well established, it is likely involved in the pathogenesis of various cancer types, as it is overexpressed in gastric cancer (Ma, Li, Zeng, Zheng, & Kang, 2023) and downregulated in bladder cancer tissues (Tan, Fu, Zuo, Wang, & Wang, 2023).

Among the other genomic risk loci that did not reach GWAS significance, only the intronic variant rs113181552 retained the association after meta-analysis with the replication cohort (**Table 1**). This variant is within the *ENSG00000286787* gene, encoding an antisense RNA that inhibits translation of the *PSMF1* gene. *PSMF1* codifies a subunit of proteasome inhibitor (also known as PI31) and has already been associated with Parkinson’s disease (Navarro & Esteras, 2024). Furthermore, its overexpression has been implicated in cancer pathogenesis by inhibiting proteasome activity and increasing cellular resistance to apoptosis induced by endoplasmic reticulum (ER) stress (Lam et al., 2023). Since altered proteasome function is implicated in numerous age-related diseases (Gomes, 2013), rs113181552 may contribute to longevity in the Calabrian population. Impaired proteostasis is indeed a hallmark of aging, whereas efficient proteostasis supports longevity (Hipp, Kasturi, & Hartl, 2019).

Overall, the association of lncRNA variants with longevity in this study may underscore the role of this class of RNAs in aging processes.

In agreement with studies carried out in Southern Italian populations, we did not find any association between the *APOE* and *FOXO3* loci and longevity in our cohort (**Table S4 of Supplementary File 1**), likely reflecting the unique genetic background of the Calabrian population (Gurinovich et al., 2019). *APOE* (apolipoprotein E) carries two missense variants that define three haplotypes − the *APOE*\*2 allele (rs429358-T/rs7412-T), the *APOE*\*3 allele (rs429358-T/rs7412-C), and the *APOE*\*4 allele (rs429358-C/rs7412-C) − and has been widely associated with longevity across diverse populations (Abondio et al., 2019; Broer et al., 2015; Deelen et al., 2019; Gurinovich et al., 2019; Sebastiani et al., 2019; Smulders & Deelen, 2024). Specifically, the *APOE*\*2 allele is generally enriched, whereas the *APOE*\*4 allele is depleted among long-lived individuals (Abondio et al., 2019; Broer et al., 2015; Deelen et al., 2019; Gurinovich et al., 2019; Sebastiani et al., 2019; Smulders & Deelen, 2024). Additionally, the variant rs7412-C has been strongly associated with reduced lifespan, highlighting the protective role of rs7412-T in age-related diseases (Satria et al., 2025). However, as mentioned, these associations were not observed in our sample as well as in similar studies carried out in Southern Italians (Gurinovich et al., 2019). The lack of *APOE* replication might be due to differences in allele frequencies worldwide and differences in population genetic backgrounds (Abondio et al., 2019; Gurinovich et al., 2019). Similarly, a recent study failed to replicate the association between *APOE* and longevity in the Sardinian Blue Zone population (Errigo, Dore, Mocci, & Pes, 2024), suggesting that population effects on this gene may be common in Southern Italy. This outcome may be explained by the high frequency of the *APOE*\*3 allele in Southern Italy (Corbo, Scacchi, Mureddu, Mulas, & Alfano, 1995) and by a latitudinal gradient in the frequency of the *APOE*\*4 allele across Europe and the Italian peninsula, as well as an ethnic-specific effect of the *APOE*\*4 allele (Abondio et al., 2019; Gurinovich et al., 2019).

Together with *APOE*, the *FOXO3* gene has been widely associated with longevity in several populations (Anselmi et al., 2009; Bae et al., 2018; Broer et al., 2015; Y. Li et al., 2009; Soerensen et al., 2010). *FOXO3* encodes the forkhead box O3 transcription factor, whose activation and repression are critical for regulating several processes, including the insulin/IGF-1 signaling pathway and the oxidative stress response (Sanese, Forte, Disciglio, Grossi, & Simone, 2019). Similarly to *APOE*, the lack of association with *FOXO3* is likely due to the genetic history of the Calabrian population, consistent with previous findings in Calabrian cohorts (Di Cianni et al., 2013). The same study also associated the glutathione S-transferase zeta 1 (*GSTZ1*) with longevity (Di Cianni et al., 2013). *GSTZ1*, highly expressed in the liver, participates in detoxification pathways and has been implicated in the liver detoxification system (Allocati, Masulli, Di Ilio, & Federici, 2018; Hayes, Flanagan, & Jowsey, 2005), hepatocellular carcinoma suppression (J. Li et al., 2019), and bladder cancer risk under arsenic exposure (Pomares-Millan et al., 2025). In our data, *GSTZ1* showed only a weak gene-based association (*p* = 0,014, **Table S6 of Supplementary File 1**). Still, we detected a genomic risk locus potentially linked to *GSTZ1* expression: the 3’-UTR variant rs11850771 in *TMED8* (Transmembrane P24 Trafficking Protein Family Member 8), which affects the expression of several genes and lies within an LD block of 43 SNPs (**Table S2 of Supplementary File 1**). Remarkably, this LD block includes the *GSTZ1* missense variant rs1046428 (p.Met82Thr, r^2^ = 0.70; **Table S3 of Supplementary File 1**) and 24 additional SNPs, all of which are eQTLs for this gene influencing its expression (**Table S3 and Table S5 of Supplementary File 1**).

Overall, beyond potential population-specific effects, our results point to biological processes already associated with aging and longevity, such as genome integrity, apoptosis, cancer pathogenesis, insulin metabolism, and inflammation (López-Otín et al., 2013, 2023). Genomic alterations, such as mutation accumulation, impaired DNA repair, and telomere shortening, are hallmarks of aging, while maintaining genome integrity is fundamental for longevity (López-Otín et al., 2013, 2023). Among the genomic risk loci identified herein, the intronic variant rs10106995, located within a lncRNA gene of unknown function (*ENSG00000300442*), may influence DNA repair by modulating *NEIL2* expression (**Table S5 of Supplementary File** 1). *NEIL2* encodes a DNA glycosylase involved in base excision repair of oxidative damage and has been linked to longevity through its role in mitochondrial DNA repair (Gredilla, Sánchez-Román, Gómez, López-Torres, & Barja, 2020). Moreover, its reduced expression and activity have been associated with several cancer types (Shinmura et al., 2016), which are often pathologies related to aging (Guo et al., 2022; López-Otín et al., 2013, 2023).

Another finding related to genome maintenance is the association of the *ACD* gene with longevity in the gene-based test (*p* = 1.15 × 10^−4^, **Figure 3** and **Table S6 of Supplementary File 1**). This gene encodes the TPP1 protein (ACD shelterin complex subunit and telomerase recruitment factor), a key component of the shelterin complex (also known as telosome), which regulates telomere length and protection (Lim & Cech, 2021). Through its interaction with POT1 (protection of telomeres protein 1), TPP1 determines the recruitment of telomerase reverse transcriptase (TERT) on telomeres, promoting telomere elongation (Liu et al., 2022), and its loss induces cell apoptosis. Dysfunction of the POT1-TPP1 complex has been associated with cancers and telomere-related syndromes characterized by premature cellular senescence (Aramburu, Plucinsky, & Skordalakes, 2020). Genetic variants in *ACD* have also been associated with telomere length, some of which are linked to longer telomeres (Burren et al., 2024), supporting *ACD* as a potential marker of longevity in our cohort.

Among the apoptosis-related signals, we identified the genomic risk locus rs75901608 (**Table 1**), along with a significant gene-based association for the *TP53BP2* (tumor protein p53 binding protein 2) gene (*p* = 2.87 × 10^−4^, **Table S6 of Supplementary File 1**). However, they respectively did not replicate and did not survive multiple-testing correction. The rs75901608 variant is located within an intron of the *SH3BP5* gene and influences its expression (**Table S5 of Supplementary File 1**). *SH3BP5* encodes for the SAB protein, which mediates JNK-dependent apoptosis in response to lipotoxicity, TNFα signaling, or ER stress (Win, Than, & Kaplowitz, 2024). However, SAB may exert context-dependent effects, as high expression has been associated with poor prognosis in acute myeloid leukemia due to reduced apoptosis (M. Li et al., 2019). *TP53BP2* encodes a p53-binding protein that enhances p53 activity and, consequently, promotes the transcription of pro-apoptotic genes; reduced expression has been reported in several cancers, contributing to impaired apoptosis and increased malignant potential (Huo et al., 2023).

Beyond apoptosis, longevity in the Calabrian cohort also appears to involve metabolic pathways related to age-associated diseases, namely diabetes and cancer, inflammation and immune response, calcium antiporter activity, and monoamine transport (**Table 2**). The most significant pathway associated with longevity in our cohort is insulin receptor substrate binding (*p*-value = 1.409 × 10^−5^), which mediates insulin signaling and glucose uptake (Johnston et al., 2003). This is consistent with the preserved glucose metabolism and low insulin resistance observed in centenarians (Paolisso et al., 2001; Vitale, Barbieri, Kamenetskaya, & Paolisso, 2017), and with their low frequency of a risk variant (T allele of rs7903146, *TCF7L2* gene) for type 2 diabetes (Garagnani et al., 2013). These results support the involvement of insulin-related pathways in longevity, not only in humans but also in other species (Deelen et al., 2013; Khan, Zou, Xiang, Chen, & Tian, 2019; Vitale et al., 2017).

In addition to metabolic pathways, inflammation and cancer also appear central to longevity in the Calabrian cohort, supporting the “*inflammaging hypothesis*”. According to this theory, aging results from maladaptive chronic inflammation in response to endogenous and exogenous stressors, while longevity reflects a more balanced inflammatory response (low production of pro-inflammatory molecules) that preserves tissue homeostasis (Franceschi et al., 2000; Fulop et al., 2023; Furman et al., 2019). Chronic low-grade inflammation contributes to age-related diseases such as cancer, diabetes, cardiovascular diseases, and neurodegenerative diseases, highlighting the importance of controlling inflammatory processes for healthy aging (Furman et al., 2019).

Finally, ion channel activity has been implicated in aging and longevity (Venkatachalam, 2022), and consistent with this, Ca^2+^ antiporter activity emerged as potentially relevant in our Calabrian cohort (*p* = 5.924 × 10^−4^). Extreme human longevity has been associated with reduced expression of genes involved in excitatory neurotransmission, where minor increases in calcium influx in neurons can extend lifespan (Zullo et al., 2019). Furthermore, the loss or reduction of activity of some calcium channels of the transient receptor potential (TRP) superfamily protects endothelial cells (Z. Li et al., 2017) and neurons against oxidative stress-induced senescence (Belrose, Xie, Gierszewski, MacDonald, & Jackson, 2012). In contrast, increased activity of inositol triphosphate receptors, which are Ca^2+^ channels located in the ER, has been related to neuronal cell death in various neurodegenerative diseases (Higo et al., 2010; Staats et al., 2016; Tang et al., 2003).

## 5. LIMITATIONS

This work presents results on genetic associations with longevity in a Calabrian population, and some limitations must be highlighted that may have influenced the detection of non-significant results.

First, the small sample size did not allow for a wider identification of genetic signals associated with longevity; therefore, further studies with a larger sample size may be required to confirm these preliminary findings.

Additionally, the high number of related individuals in our cohort could lead to population structure, potentially altering the detection of genetic associations. To reduce this bias, we used PC-AiR and PC-Relate results from the GENESIS R package to account for population structure and relatedness. We incorporated this information into a mixed-model analysis to yield more reliable association results.

## 6. CONCLUSIONS

In conclusion, our results suggest that longevity in the Calabrian population exhibits a population-specific effect and is influenced by molecular mechanisms previously identified in other studies. We detected new genomic risk loci associated with this trait, among which only the rs113181552 variant was replicated and may serve as a promising marker of longevity. The absence of associations with *APOE* and *FOXO3* likely reflects the unique genetic history of this population, despite their known effects in other cohorts. Overall, this study supports the involvement of key mechanisms underlying longevity, including protein homeostasis, DNA repair and telomere maintenance, apoptosis, insulin and cancer-related pathways, calcium antiport, and inflammation.

In the future, it will be interesting to evaluate dominant and recessive genetic effects on this trait and to test rare variants and the X chromosome in association with longevity. It will also be valuable to deepen the genetic history of this population, since all individuals showed a percentage of Ashkenazi Jewish ancestry, and to verify if there are signs of natural selection in the genome that may be related to longevity in this population.

## AUTHOR’S CONTRIBUTION

Idea/concept: LB, GR, SD, VN; Data curation/processing: LB, PC, BTB, SD; Data analysis: LB; Writing: LB, GR, SD, VN; Advice/review: SP, JH, MEB, GP.

## AKNOWLEDGMENTS

Funding: The European Union funded this research (to V.N. and S.D) – Next Generation EU, through the Italian Ministry of University and Research, within the framework of the PRIN 2022 project ‘EXTREMAL: EXTREMe phenotypes for Aging and Longevity’ [D.D.104 2.02.2022; cod. 20227KNBEJ, PNRR M4.C2.1.1].

## CONFLICTS OF INTEREST

The authors declare no conflict of interest.

## DATA AVAILABILITY

GWAS summary statistics will be uploaded to the GWAS catalog.

## Notes

### Competing Interest Statement

The authors have declared no competing interest.

