## Supplementary Figures for "Genetic Associations with Longevity in a Calabrian Cohort: A Genome-Wide Study"

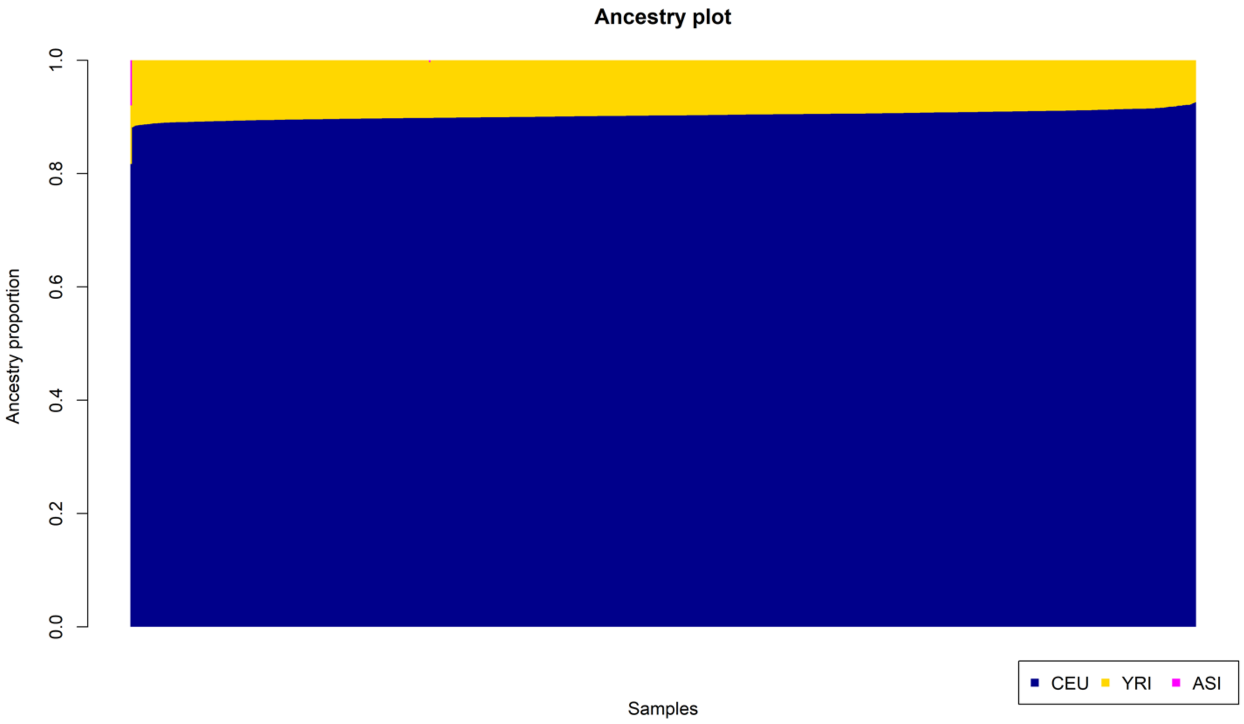


**Supplementary Figure 1.** Ancestry plot of Calabrian individuals included in the study, evaluated using a panel of three different populations: Northern Europeans from Utah (CEU), Yoruba population (YRI), and East Asian populations (ASI, comprehending Han Chinese in Beijing (CHB) and Chinese in Metropolitan Denver (CHD)). All individuals show European ancestry (CEU) ranging from 81 – 92%, and minor African ancestry (YRI) ranging from 7 – 10%. Only one individual shows small portion (7%) of Asian ancestry (ASI).

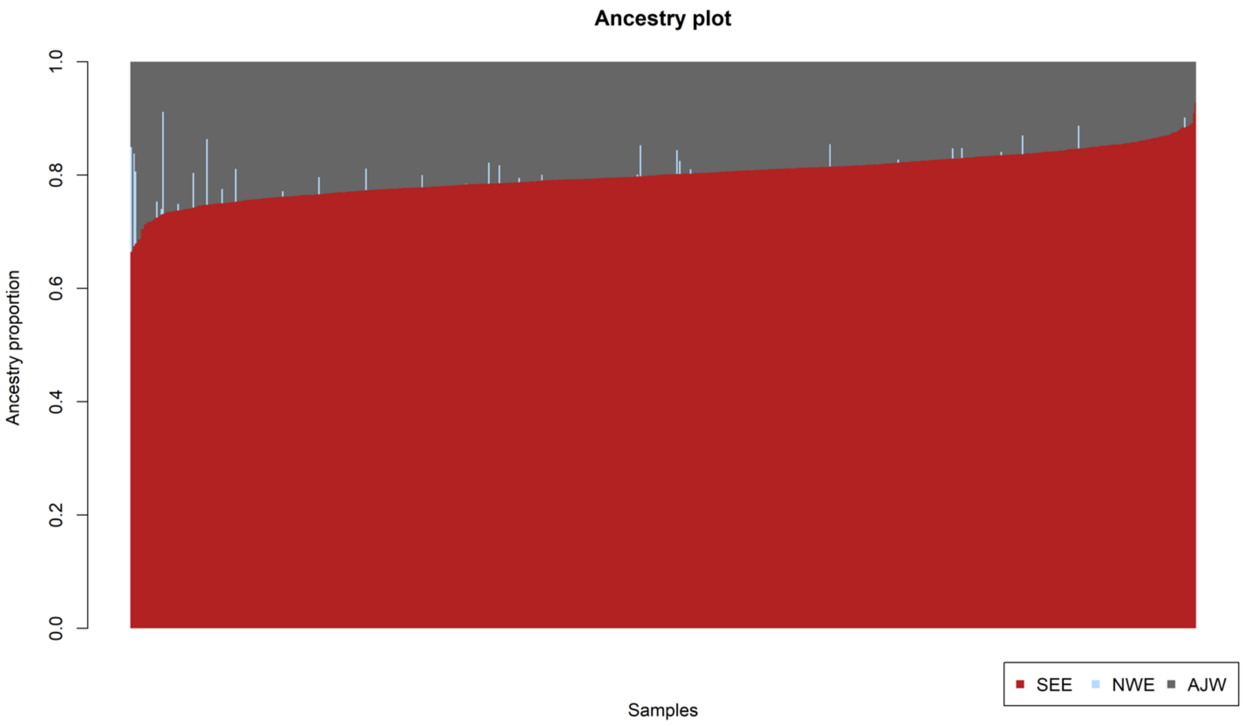


**Supplementary Figure 2.** Ancestry plot of Calabrian individuals included in the study, evaluated using a panel of three different populations of European ancestry: Southeastern European (SEE), Northwestern European (NWE), and Ashkenazi Jews (AJW). Southeastern European is the prevalent ancestry is the Southeastern European (66 – 93 %), followed by Ashkenazi Jews (7 – 33 %). 33 individuals display a percentage of Northwestern European ancestry (0.1 – 18 %).
